# Immunoregulatory macrophages modify local pulmonary immunity and ameliorate hypoxic-pulmonary hypertension

**DOI:** 10.1101/2023.07.31.551394

**Authors:** Angeles Fernandez-Gonzalez, Amit Mukhia, Janhavi Nadkarni, Gareth R. Willis, Monica Reis, Kristjan Zhumka, Sally Vitali, Xianlan Liu, Alexandra Galls, S. Alex Mitsialis, Stella Kourembanas

## Abstract

**Rationale:** Macrophages play a central role in the onset and progression of vascular disease in pulmonary hypertension (PH) and cell-based immunotherapies aimed at treating vascular remodeling are lacking.

**Objective:** To evaluate the effect of pulmonary administration of macrophages modified to have an anti-inflammatory/pro-resolving phenotype in attenuating early pulmonary inflammation and progression of experimentally induced PH.

**Methods:** Mouse bone marrow derived macrophages (BMDMs) were polarized *in vitro* to a regulatory (M2_reg_) phenotype. M2_reg_ profile and anti-inflammatory capacity were assessed *in vitro* upon lipopolysaccharide (LPS)/interferon-γ (IFNγ) restimulation, before their administration to 8- to 12-week-old mice. M2_reg_ protective effect was tested at early (2 to 4 days) and late (4 weeks) time points during hypoxia (8.5% O_2_) exposure. Levels of inflammatory markers were quantified in alveolar macrophages and whole lung, while PH development was ascertained by right ventricular systolic pressure (RSVP) and right ventricular hypertrophy (RVH) measurements. Bronchoalveolar lavage (BAL) from M2_reg_-transplanted hypoxic mice was collected, and its inflammatory potential tested on naïve BMDMs.

**Results:** M2_reg_ macrophages demonstrated a stable anti-inflammatory phenotype upon a subsequent pro-inflammatory stimulus by maintaining the expression of specific anti-inflammatory markers (Tgfß, Il10 and Cd206) and downregulating the induction of proinflammatory cytokines and surface molecules (Cd86, Il6 and Tnfα). A single dose of M2_regs_ attenuated the hypoxic monocytic recruitment and perivascular inflammation. Early hypoxic lung and alveolar macrophage inflammation leading to PH development was significantly reduced and, importantly, M2_regs_ attenuated RVH, RVSP and vascular remodeling at 4 weeks post treatment.

**Conclusions:** Adoptive transfer of M2_regs_ halts the recruitment of monocytes and modifies the hypoxic lung microenvironment, potentially changing the immunoreactivity of recruited macrophages and restoring normal immune functionality of the lung. These findings provide new mechanistic insights on the diverse role of macrophage phenotype on lung vascular homeostasis that can be explored as novel therapeutic targets.

## INTRODUCTION

Inflammation and immune dysregulation are known factors contributing to the pathogenesis of pulmonary arterial hypertension (PAH), a condition characterized by progressive vascular remodeling and increased pulmonary vascular resistance [1, 2]. Strong accumulating data indicates that monocytes and macrophages play a critical role in the induction and progression of PAH. For example, CD68^+^ macrophages are predominant in advanced obliterative plexiform lesions from PAH patients [3–5] and pulmonary arterial adventitia of chronically hypoxic models of pulmonary hypertension [6, 7] (by convention referred as having PH). Macrophages in the lung are heterogeneous cell populations occupying different niches and exhibiting microenvironment specific phenotypes and functions. Attempts to eliminate discrete monocyte and macrophage subtypes or modify their immune response have been explored in experimental PH [8–11]. However, monocyte/macrophage depletion by pharmacological and genetic approaches have yielded conflicting results regarding the role of myeloid cells in regulating PH pathogenesis [9, 12, 13]. Macrophage activation and cytokine production are also potential targets in the progression of PH. Using a murine model of hypoxia-induced PH, our group reported that alternatively activated macrophages contribute to the development of vascular remodeling demonstrating that an early “alternative” macrophage activation and upregulation of inflammatory markers is essential for vascular remodeling and establishment of PH [14]. Concurrently, macrophage activation can also be driven by adventitial fibroblasts through paracrine interleukin 6 (Il6) signaling [15] and by activation of the Il6/Il21 signaling axis in T-helper-17(Th17) cells [16]. Anti-inflammatory therapies aimed to inactivate or eliminate the occurrence of “abnormally” activated macrophages in these studies have targeted diverse cellular components of the alveolar and interstitial lung compartments and have led to a new interest in lung microenvironmental signals and their producing cells in the search of medicinal alternatives. Adoptive-transfer and transplantation techniques employing bone-marrow-derived and pulmonary macrophages have been successfully investigated as strategies for increasing the number of functional macrophages [17]. Macrophages modified to have an anti-inflammatory phenotype have been used to ameliorate inflammation, pathological structural changes, and functional decline during kidney injury [18], spinal cord injury [19], colitis-induced mucosal inflammation [20] organ transplantation [21] and type 1 diabetes [22].

As a novel approach, we evaluated the impact of pulmonary administration of anti-inflammatory/pro-resolving macrophages, polarized *in vitro* with IL10, TGFβ and IL4 cytokines, in restoring lung immune homeostasis, attenuating early pulmonary inflammation, and ameliorating vascular injury in chronic hypoxia-induced PH with the goal to provide a platform for the application of cell-based-macrophage therapies for the treatment of PH.

## METHODS

### Animals and hypoxic exposures

Eight to 10-week-old male FVB mice were used for most experiments in this study. Heme oxygenase1 null mice (*Hmox1^−/−^*), created in a mixed 129vJ × Balb/c background were bred in our facility and were used at 8 - to 10-weeks of age for hypoxia experiments. The hypoxia-induced PH mouse model has been well-established and used by our group in previously published work [14, 23, 24]. Briefly, 8- to 10-week-old mice were housed inside a Plexiglass chamber (BioSpherix, Parish, NY) where normobaric hypoxia (8.5 ± 0.5 % O_2_) was induced by an Oxycycler controller (Biospherix, Ltd., Lacona, NY). Levels of CO_2_ were maintained at <0.5% (less than 5,000ppm) and ammonia was removed by ventilation and activated charcoal filtration using an electric air purifier. All procedures were in accordance with regulations approved by the Boston Children’s Hospital Animal Care and Use Committees (IACUC).

### Culture and polarization of bone marrow derived macrophages

Macrophages were isolated from femurs and tibiae of 5-to 7-week-old FVB mice. Mouse bone marrow cells were flushed with DMEM media. The cell suspension was cultured in non-treated culture plates for 7 days to obtain bone marrow-derived macrophages (BMDMs), using DMEM medium containing 10% fetal bovine serum (FBS), penicillin (100U/ml), streptomycin (100μg/ml), and recombinant murine M-CSF (Peprotech, Rocky Hill, NJ) or L929-cell conditioned medium as described previously [25]. Media was supplemented every 2-3 days. The cells cultured for 6-7 days in the above medium were defined as naïve macrophages. For generation of immunoregulatory macrophages (hereinafter M2_regs_), naïve macrophages were incubated for 24h with 20ng/ml TGFβ, 20ng/ml IL10, and 20ng/ml IL4 (Peprotech, Rocky Hill, NJ). For repolarizarion, M2_regs_ and untreated (M0, cultured in medium alone) cells were washed and treated with 20ng/ml lipopolysaccharide (LPS) and 20ng/ml interferon-γ (IFNγ) for another twenty-four hours.

### Flow cytometry

For characterization of BMDMs and myeloid populations prepared from digested lungs at two-days and 4-weeks post-hypoxia, cells were incubated with the labeling antibodies listed in Table S1. Briefly, lung was digested with a mixture of Collagenase IV (1.6 mg/ml) and DNase (50 unit/ml) (Worthington) at 37°C for 1 h. Erythrocytes were removed with lysis buffer (Roche) and the remaining cell suspension filtered through a 40μm nylon mesh (Corning, MA, US). Cell flow was pelleted, washed, and counted using an automated cell counter. Cells were stained with the appropriate fluorochrome-conjugated antibodies. The gating strategy for flow cytometry to characterize *in vitro* macrophage polarization and analysis of *ex vivo* macrophages recovered from lung tissue are shown in figures S1 and S2. Data were acquired on a LSRFortessa cell analyzer (BD Biosciences, MA, US). Compensation was adjusted accordingly and supported by UltraComp ebeads (Affymetrix, CA, US). Cell populations were identified using sequential gating strategy and recorded as a percentage of the total cell population. The expression of macrophage activation markers is presented as median fluorescence intensity (MFI), calculated post-acquisition using FlowJo 10.8 (BD Biosciences) software.

For *in vivo* cell tracking, cells were labeled with DiI (1,1’-Dioctadecyl-3,3,3’,3’-tetramethylindocarbocyanine perchlorate) or DiD (1,1’-Dioctadecyl-3,3,3’,3’-tetramethylindocarbocyanine 4-chlorobenzenesulfonate salt) dye for 20 min at 37°C. The labeling was stopped by washing the cells twice with warm DMEM medium containing 10% Fetal Bovine Serum. Cells were recounted and suspended at a concentration of 0.6 to 1 × 10^6^ BMDMs in 25μl (intranasal) or 50μl (endotracheal) of DMEM media. Cells were administered to anesthetized FVB recipients. Control animals received an equivalent volume of DMEM media only. After transfer, mice were rested for 2 to 4 hours and then exposed to hypoxia for 2 (to analyze lung myeloid cell populations) or three consecutive days (to investigate inflammatory pathways) or four consecutive weeks (to examine pulmonary and cardiac structural changes), in a Plexiglass chamber with full access to food and water. The analysis of DiI-labeled spatial distribution by both oropharyngeal and intranasal routes was assessed in lung sections obtained from mice after three days and 4 weeks post-transplantation. The detection of DiI immunofluorescence staining (Figure S3A, top panel) determined both routes were efficacious for macrophage intrapulmonary localization and were used interchangeably in subsequent experiments. Moreover, bronchoalveolar (BAL) lavage of lungs allowed the retrieval of DiI-stained macrophages and their detection in cytospins prepared from normoxic (Nrmx) and hypoxic (Hpx) mice (Figure S3B) and indicated M2_regs_ accumulate preferentially in the alveolar space (Figure S3A, mid panel).

### Bronchoalveolar lavage

Mice were anesthetized with pentobarbital (75mg/kg i.p.) and their trachea cannulated with a blunt ended 20gauge Luer Stub Adapter (Becton Dickinson). BAL fluid (BALF) was collected via sequential administration of 6-8 lavages with 0.5ml Hank’s Balanced Salt solution (HBBS) supplemented with 10 mM EDTA and 1mM HEPES and approximately a volume of 3.5-4.0 ml of BALF was recovered per mouse. Cells in BALF were collected by centrifugation at 400 *x g* for 7 min and leukocytes counted with an automated cell counter.

### Right ventricular systolic pressure and right ventricular weight measurements

After four weeks in hypoxia, mice were anesthetized and right ventricular systolic pressure (RVSP) was measured using a closed chest approach and the PowerLab system (ADInstruments, Colorado Springs, CO), as previously described [26]. Mice were then sacrificed with an overdose of pentobarbital, and lungs were inflated and fixed with 4% paraformaldehyde after perfusion with phosphate buffered saline (PBS). Lung samples were paraffin-embedded and sectioned and heart samples were used for Fulton’s index measurements (ratio between right ventricular weight and left ventricle plus septal weight, RV/[LV+S]), an assessment of RVH.

### Lung Histology and Immunofluorescence

Paraffin lung sections were rehydrated and subjected to antigen unmasking with a 10mM citrate buffer. Cytospin preparations obtained from BALF were fixed in 4% paraformaldehyde. Both tissue sections and cytospins were incubated in blocking buffer containing 4% normal serum and incubated with primary antibodies (Table S1) against ⍺-smooth muscle actin (αSMA), von Willebrand factor (vWf), found in inflammatory zone 1 (Fizz1), the macrophage marker CD68 and complement C5a receptor 1 (C5aR1). Following overnight incubation, slides were washed and incubated with Alexa Fluor^TM^ 488 or 555-conjugated secondary Abs, counterstained with 4’,6-diamino-2-phenylindole (DAPI) to visualize the nuclei, dried and visualized under fluorescence microscopy using a Nikon Eclipse 80i microscope (Nikon, Tokyo, Japan). Pulmonary vascular remodeling was determined by quantifying the vessel wall thickness and the percentage of muscularized vessels using Metamorph software v.6.2.r (Universal Imaging, Downingtown, PA). The vessel wall thickness was measured in 50-150μm ⍺SMA-stained sections captured at 200x magnification, using the following equation: Medial Thickness Index = [(area_ext_ − area_int_)/area_ext_], where area_ext_ and area_int_ are the areas within the external and internal boundaries of the αSMA layer, respectively. The percentage of muscularized vessels was quantified in vWf- and αSMA double-stained non-overlapping images at 100× magnification by counting the number of vWf-positive peripheral vessels (20-100μm) stratified by αSMA content. These vessels were classified as non-muscular (NM), partially muscular (PM), or fully muscular (FM), according to αSMA staining. Vessels that were αSMA positive but lacked a continuous layer were considered PM while FM vessels had a continuous αSMA band.

### Reverse transcription-polymerase chain reaction analysis

Total lung, alveolar macrophages from BAL collection, and BMDM mRNA isolation and analysis was performed by RT-qPCR as previously described (Willis et al., AJRCCM 2018). Briefly, TRIzol (ThermoFisher) was used to extract total RNA, as per manufacturer’s instructions. Taqman® primers listed in Table S2 were used in the PCR reactions. Nucleoporin 133 (Nup133) served as an internal standard. Annealing was conducted at 60°C for 30 sec, extension at 72°C for 30 sec, and denaturation at 95°C for 30 sec for forty cycles in an StepOnePlus Thermocycler (Applied Biosystems, Waltham, MA).

### Statistical analysis

Data were analyzed with GraphPad Prism 9 (GraphPad software, San Diego, CA) using one-way ANOVA followed by Tukey’s or Sidak’s multiple comparison test between groups. Differences between two groups were compared using Mann-Whitney test. Values represent the mean ± SEM unless otherwise indicated. Statistical differences are denoted as *p < 0.05, **p < 0.01, ***p < 0.001, ****p <0.0001.

## RESULTS

### Polarized IL4/IL10/TGF**β** macrophages adopt and retain an anti-inflammatory phenotype *in vitro*

BMDMs polarized with IL4/IL10/TGFβ for 24 hr showed induced mRNA expression of M2-associated markers, namely *Tgfβ*, *Il10*, *Arg1*, *Cd206 (Mrc1)* and *Pdl2* and reduced mRNA levels of *Cd86*, a surface marker associated with macrophage M1 activation (Figure 1A). In agreement with the gene expression, CD86 surface levels in M2_regs_ were comparable to that in untreated (M0) macrophages while CD206 and PDL2 surface expression was increased above untreated controls (Figure 1B). Given that chronic hypoxia induces an inflammatory condition that may restrict/restrain the protective effect of macrophage adoptive transfer, we investigated whether a secondary pro-inflammatory stimulation would impair the anti-inflammatory phenotype of M2_reg_ using an *in vitro* experimental approach (Figure1C). To that end, we compared the response of non-treated cells with the response of established M2_regs_ to restimulation with lipopolysaccharide (LPSγ/IFNγ) treatment. Compared to untreated cells (N-LPS/IFNγ), M2_regs_ displayed increased mRNA expression of *Tgfβ*, *Il10* and *Cd206* and reduced mRNA levels of the typical M1 markers *Cd86*, *Il6* and *Tnf⍺* when the cells were washed and stimulated with LPS/IFNγ for 24 hrs (Figure 1C). Accordingly, the M2 surface marker CD206 expression was higher while CD86 expression was still significantly lower in M2_regs_ compared to untreated cells, following restimulation (Fig. 1D). In summary, these data indicate that activation in M2_regs_ leads to an anti-inflammatory phenotype that prevents subsequent M1 repolarization *in vitro*.

**Figure 1.**
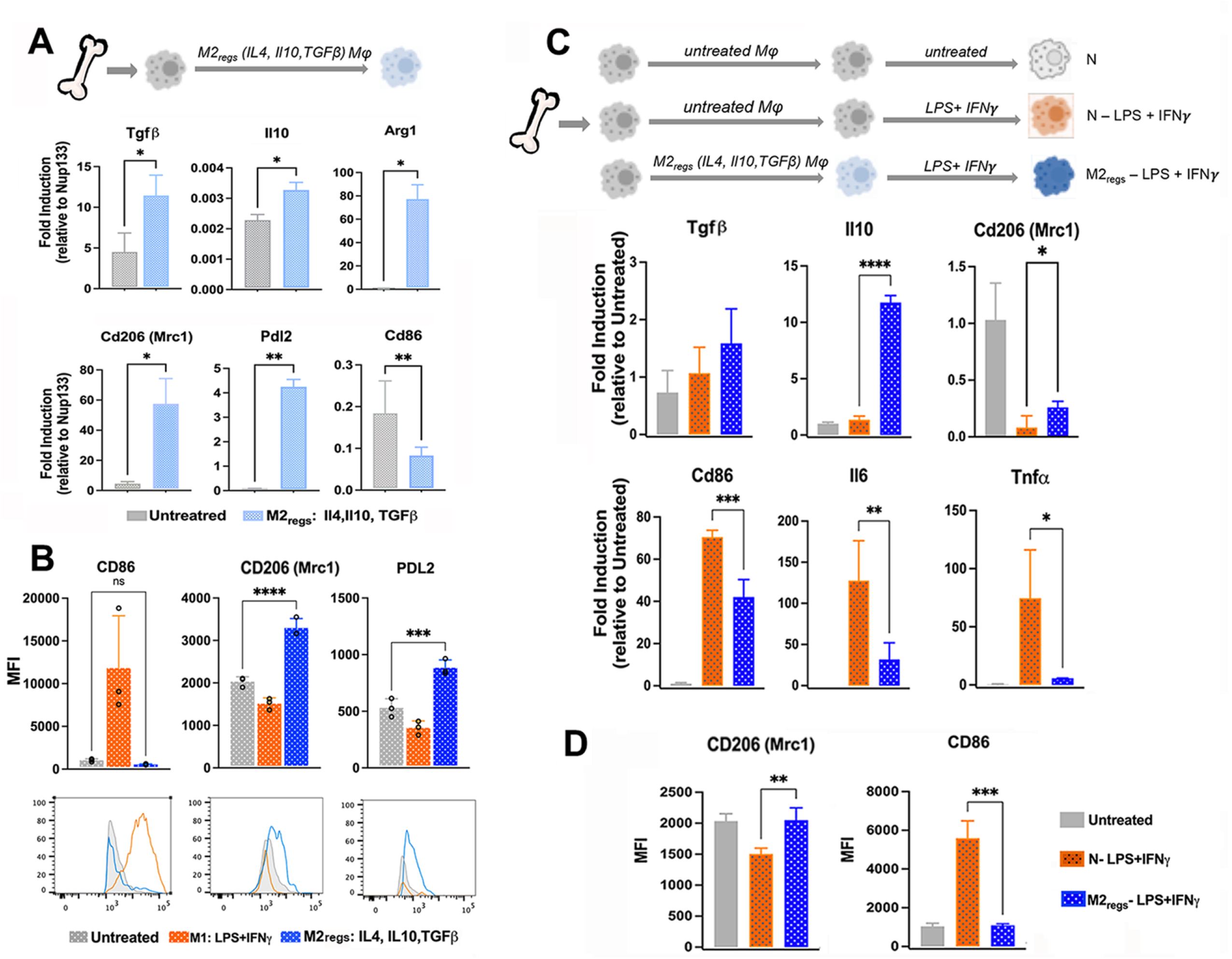
Established M2regs show reduced pro-inflammatory phenotype after restimulation with LPS and IFNγ. (A) M2_regs_ (IL4, IL10, TGFβ) showed induced mRNA expression of alternatively (M2) associated markers *Tgfβ*, *Il10*, *Arg1*, *Cd206* and *Pdl2* and reduced mRNA levels of *Cd86*, a proinflammatory surface marker associated with macrophage activation and M1 activation compared to M0 (untreated) cells. (B) Flow cytometry was used to assess CD86, CD206 and PDL2 surface expression (median fluorescence intensity, MFI) in M2_reg_ and M1 (LPS/IFNγ) polarized BMDMs (used here as controls). Representative histograms for CD86, CD206 and PDL2 MFIs are shown. (C) Restimulated M2_regs_ retain their M2-associated phenotype (increased *Tgfβ*, *Il10* and *Cd206* mRNA) and reduced expression of M1/classical markers (*Cd86, Il6 and Tnf⍺*) in response to LPSγ/IFNγ, compared with non-treated (N-LPS/IFNγ) cells. (D) CD206 and CD86 surface expression (MFI) in restimulated M2_regs_-LPS/IFNγ assessed by flow cytometry. Results are representative of 2-4 independent experiments. Mean ± SD, n=3-5 per group, *p <0.05, **p <0.01, ***p <0.001, ****p <0.0001.

### Adoptive transfer of M2_regs_ attenuates chronic hypoxia-induced PH

To determine if M2_reg_ macrophages attenuated the development of hypoxia-induced PH and the degree of associated pulmonary vascular remodeling, a single dose of 0.6-1 × 10^6^ M2_regs_ was adoptively transferred to 8- to 10-week-old mice as shown in Fig. 2A. Four weeks after the delivery of M2_regs_, Nrmx and Hpx mice were sacrificed and indexes of PH such as right ventricular (RV) hypertrophy (right ventricular weight/left ventricular plus septum weight, RV/LV+S), RV systolic pressure (RVSP) and vascular remodeling (Medial Thickness Index and vessel muscularization) were evaluated.

**Figure 2.**
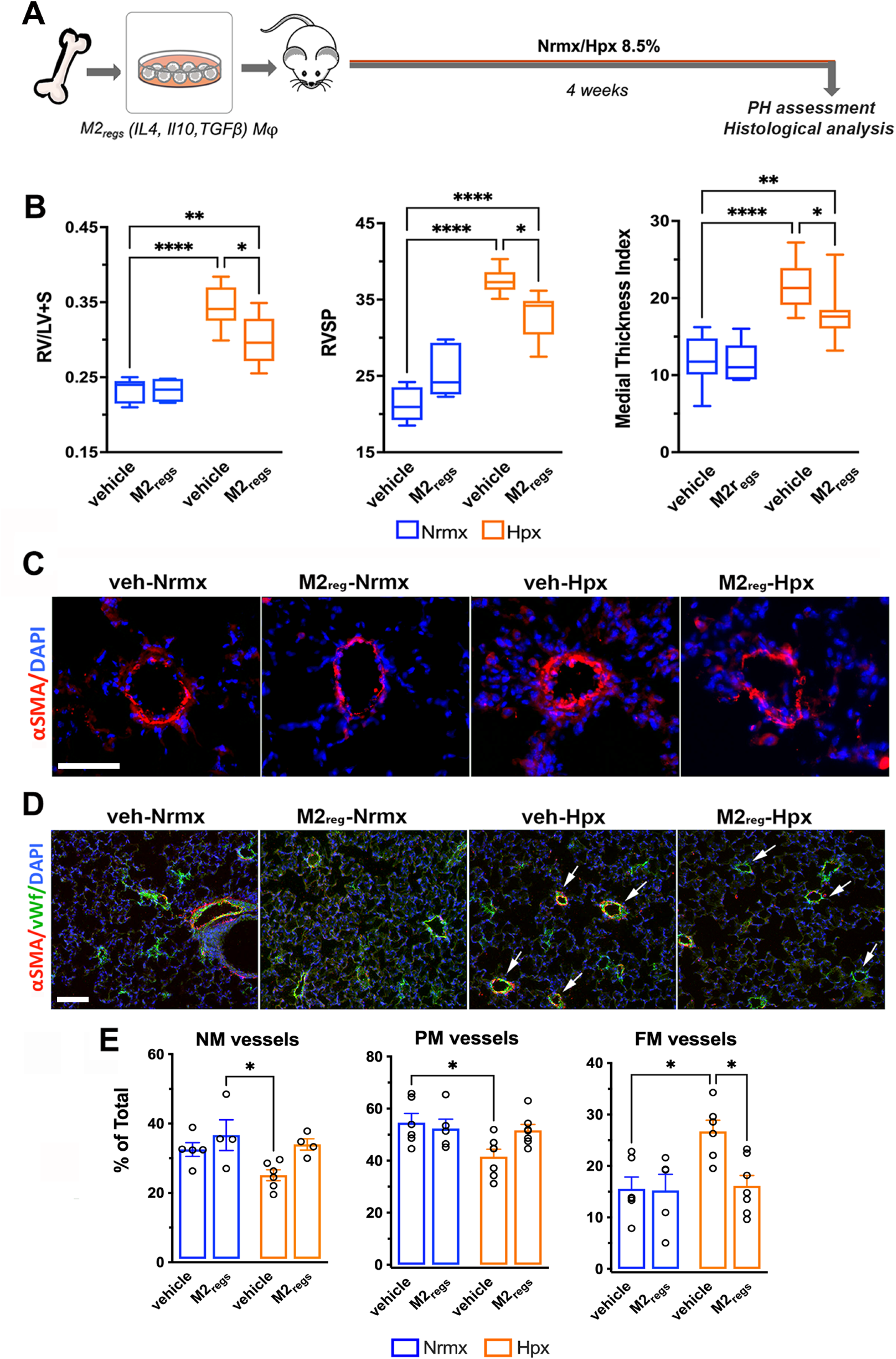
Adoptive transfer of M2_regs_ reduces vascular injury and PH progression. (A) Mice received 0.6-1 × 10^6^ M2_regs_ or media (vehicle) by intrapulmonary administration followed by exposure to normal air (Nrmx) or hypoxia (8.5% oxygen, Hpx) for 4 weeks. (B) Fulton’s Index (RV/LV+S), right ventricular systolic pressure (RSVP) and Medial Thickness Index determined after long-term exposure. (C) Representative images of lung vascular remodeling in small (50-150μm) pulmonary arterioles (PA) of vehicle (veh) and M2_regs_-treated Nrmx or Hpx mice, as assessed by α-smooth muscle (αSMA) staining; scale bar = 50μm. (D) Peripheral muscularization of PAs in the lungs of Hpx mice and effect of M2_reg_ administration, analyzed by von Willebrand factor (vWf) and αSMA double immunofluorescence; arrows indicate colocalization in small (20-100μm) PAs; scale bar = 50μm. (E) Quantification of muscularization of non-muscular (NM), partially muscular (PM), and fully muscular (FM) PAs is shown in D. Mean ± SEM, n=4-6 per group, *p <0.05, **p <0.01, ****p <0.0001.

In vehicle treated mice, exposure to chronic Hpx induced an increase in Fulton’s index and RVSP. However, a single administration of M2_regs_ attenuated both RV hypertrophy and RVSP (0.299 ± 0.015 vs 0.344 ± 0.012, p<0.05 and 32.820 ± 1.047 vs 37.51 ± 0.668, p<0.05, respectively). Furthermore, distal pulmonary arterial remodeling as assessed by ⍺SMA staining was less marked in M2_reg_-treated Hpx mice compared to vehicle counterparts (17.77 ± 1.038 vs 21.52 ± 0.928, respectively, p<0.05) (Fig. 2B, C). Additional analysis of peripheral vessel muscularization of by dual αSMA and vWf immunofluorescence staining revealed that hypoxia increased the number of fully-muscularized (FM) small arteries in the lung at the expense of non-muscularized (NM) and partially muscularized (PM) vessels, and this effect was prevented by M2_regs_ administration (Fig. 2D, E). These data suggest that adoptive transfer of M2_regs_ has a protective effect on the development of PH induced by chronic hypoxia exposure.

To test the retention of the anti-inflammatory phenotype of M2_regs_ *in vivo*, DiD-labeled M0 and M2_regs_ were adoptively transferred to mice, which next remained in Nrmx or were exposed to Hpx (Figure 3A), and the *ex vivo* surface expression of CD45, CD11c, SiglecF, CD86, CD206 and PDL2 of retrieved M0 and M2_regs_ determined by flow cytometry. Expression of CD86 and PDL2 by M2_regs_ has been previously well characterized by Parsa and colleagues, in a murine model of type I diabetes where they demonstrated that stimulation with IL4/IL10/TGF generate macrophages that retain an anti-inflammatory phenotype *in vitro* as well as *in vivo*, and the ability to suppress T-cell proliferation [22]. In our study, the frequency of CD86^+^-M2_regs_ was reduced while that of M2_regs_ expressing CD206 and PDL2 was induced compared to untreated (M0) and LPS/IFγ (M1) BMDMs before adoptive transfer (Figure 3B). Four weeks after transfer, the percentage of labeled CD45^+^ cells found in M0 and M2_reg_ Hpx-treated lungs was reduced compared with their Nrmx littermates (56 ± 1.46 vs 70.19 ± 1.04, p <0.0001 and 55.90 ± 1.69 vs 69.28 ± 1.66, respectively, p <0.0001). In addition, the percentage of retained DiD M2_regs_ in the lung was reduced (albeit not statistically significant), when compared to the percentage of retained DiD M0s (2.8 ± 0.78 vs 4.87 ± 0.32 and 2.34 ± 1.15 vs 5.53 ± 1.50 in Nrmx and Hpx, respectively). Notably, most labeled DiD^+^ cells were CD11c^+^SiglecF^+^ at the time of analysis, characteristic markers of resident alveolar macrophages. Analysis of retained CD45^+^DiD^+^ macrophages revealed that 56.30 ± 10.9% M2_regs_ retrieved from Hpx mice, were CD86^+^ vs 38.45 ± 7.6% M0s when compared to Nrmx-M2_regs_ and M0s respectively, while the proportion of Hpx-M2_regs_ expressing PDL2 represent twice the number of Hpx-M0 (18.92 ± 5.5 in Hpx-M2_regs_ vs 8.93 ± 2.3 in Hpx-M0) when compared to Nrmx (p <0.05). Percentages of retrieved CD206^+^ DiD-M0 and M2_reg_ labeled populations were not affected by environmental exposure (3.55 ± 0.6 vs 5. 07 ± 0.7 in Nrmx and 3.46 ± 0.7 vs 5.03 ± 0.8 in Hpx) (Figure 3C). These results indicate that adoptively transferred M2_regs_ are retained in the lung, where they may adopt phenotypic characteristics of alveolar macrophages, akin to their local residence in the airspace. Furthermore, they suggest that the phenotype of polarized M2_regs_ macrophages can, to some extent be reversed *in vivo*, especially after sustained chronic inflammation.

**Figure 3.**
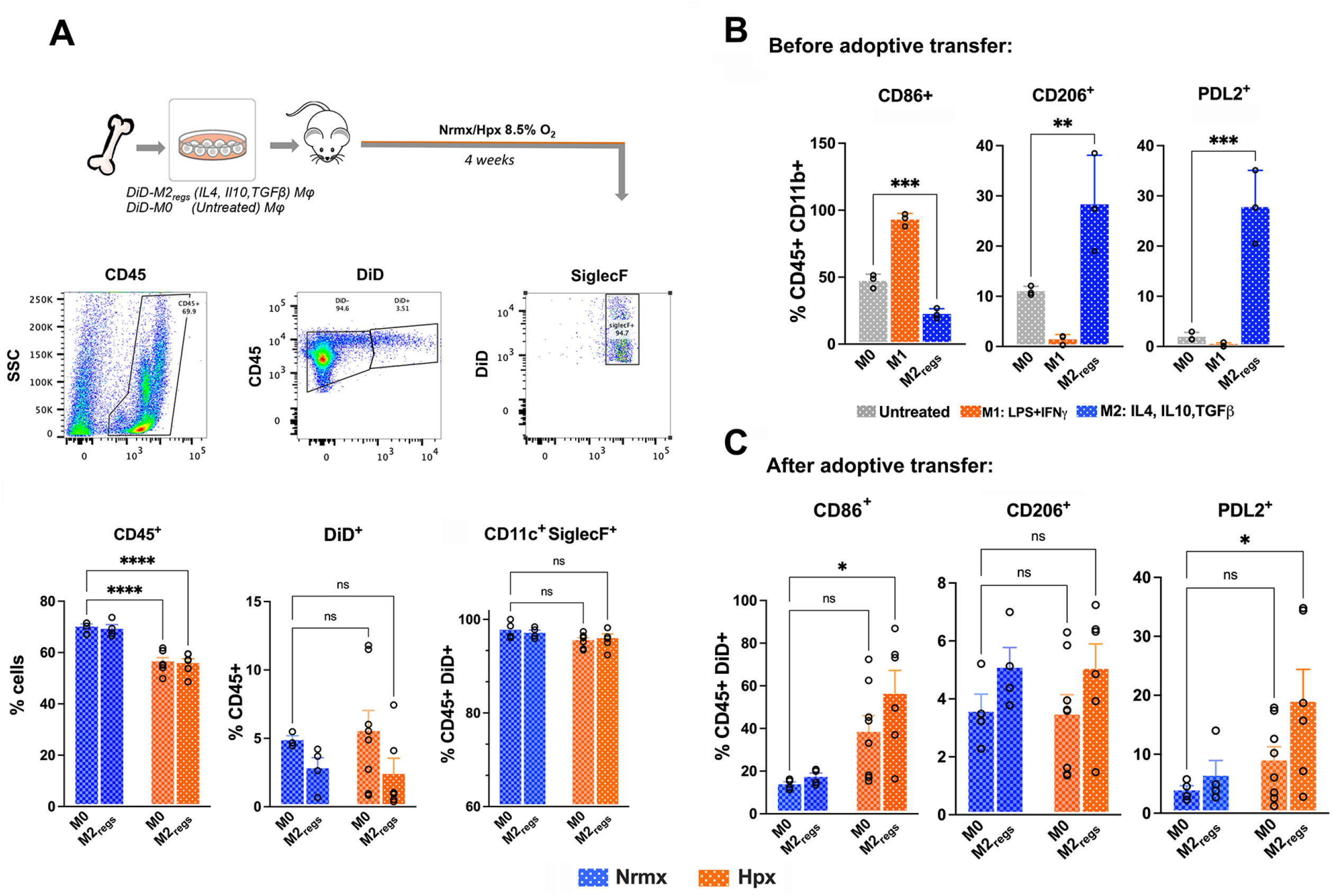
Chronic inflammation partially reverses the phenotype of M2_regs_ *in vivo*. (A) Mice received 0.6-1 × 10^6^ DiD-M2_regs_ or DiD-M0s_s_ by intrapulmonary administration followed by exposure to normal air (Nrmx) or hypoxia (8.5% oxygen, Hpx) for 4 weeks. Flow cytometry analysis of CD45^+^, DiD^+^, CD11c^+^SiglecF^+^ cells recovered from cell suspensions of Nrmx or Hpx lungs after adoptive transfer. Histograms show percentages of parental populations. (B) Representative histograms of CD86^+^, CD206^+^ and PDL2^+^frequencies showing efficient M2_reg_ polarization before their transfer to mice and exposure to chronic hypoxia. Frequencies of LPS/IFNγ stimulated cells (M1) are shown for comparison. Results are representative of three independent experiments. (B). (C) Quantification of CD86^+^, CD206^+^ and PDL2^+^ cell frequencies of CD45 DiD^+^ retrieved cells from M0 and M2_reg_-treated mice after exposure to Nrmx or Hpx. Mean ± SD, n=4-8 per group, *p <0.05, **p <0.01, ***p <0.001, ****p <0.0001.

### Adoptive transfer of M2_regs_ suppresses hypoxia-induced inflammation in lung and alveolar macrophages

To evaluate the possibility that treatment with anti-inflammatory M2_regs_ regulates the early inflammatory response leading to the progression of vascular disease, levels of inflammatory markers Il6, Ccl17, Fizz1 and Arg1 were quantified by qPCR in alveolar macrophages and whole lung after three days of hypoxia (Fig. 4A). Lungs from vehicle-treated mice had increased mRNA expression of *Il6*, *Ccl17*, and *Fizz1*. Adoptive transfer of M2_regs_ reduced the hypoxic induction of *Il6*, *Ccl17* and *Fizz1* while boosting the levels of *Arg1* compared to those seen in vehicle-treated, Hpx-exposed mice (Fig. 4B). Immunofluorescence analysis of Fizz1 in lung sections revealed increased staining in alveolar macrophages of vehicle treated Hpx-exposed mice that was absent from the lungs of M2_regs_-receiving mice (Fig. 4C). Similarly, levels of proinflammatory *Il6* and *Ccl17* mRNA, and Fizz1 protein expression quantified in BAL macrophages were increased after 3 days of Hpx, while those increases were attenuated by M2_regs_ administration (Fig. 4D, E). Collectively, these results suggest that a single administration of M2_regs_ ameliorates early hypoxia-induced inflammation leading to PH development.

**Figure 4.**
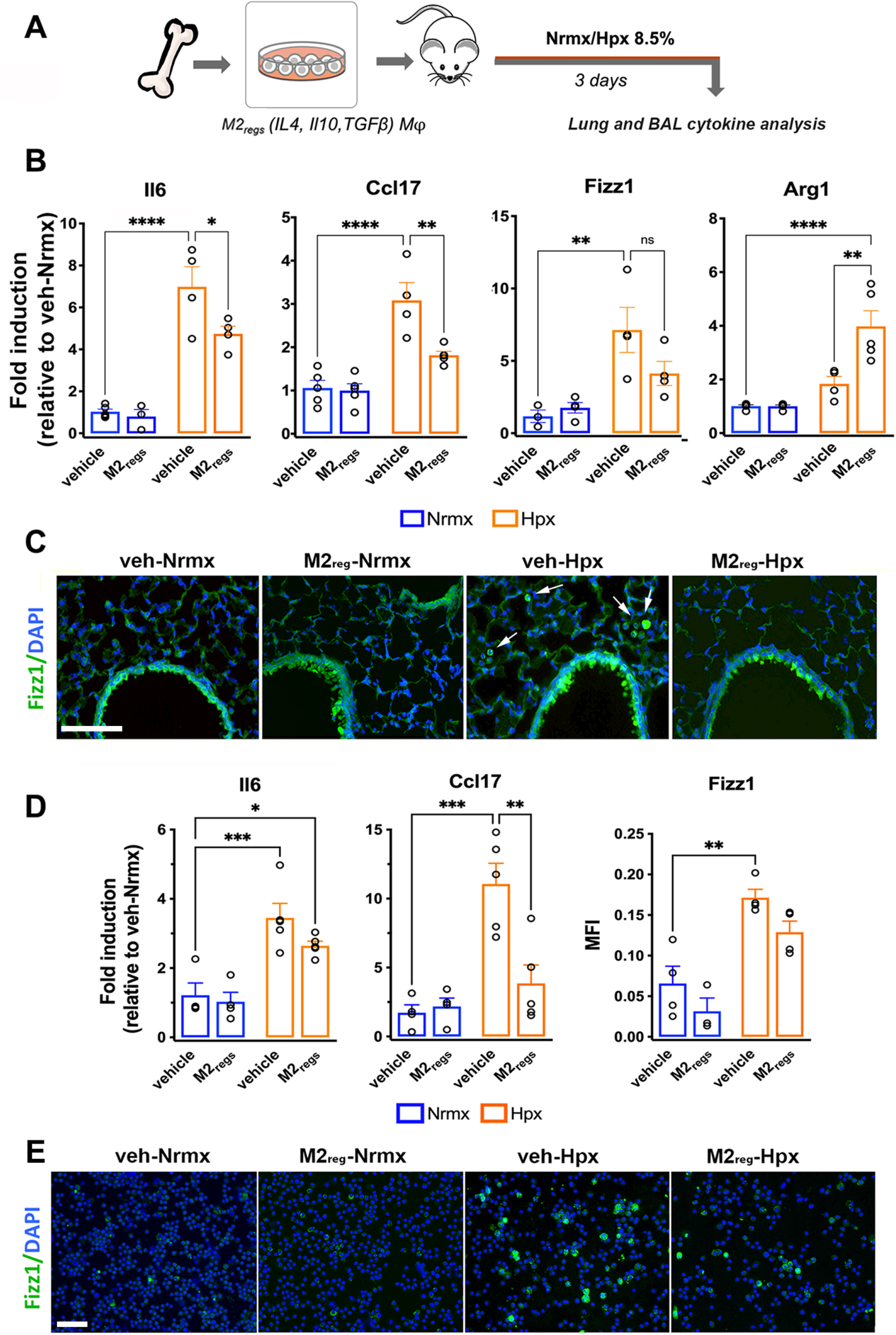
M2_reg_ adoptive transfer decreases early Hpx-induced cytokine expression in lung and BAL. (A) Mice received 0.6-1 × 10^6^ M2_regs_ or media (vehicle) by intrapulmonary administration followed by exposure to normal air (Nrmx) or hypoxia (8.5% oxygen, Hpx) for 3 days. (B) Whole lung *Il6, Ccl17, Fizz1* and *Arg1* mRNA levels were analyzed by qPCR at this time point. Expression levels are shown relative to vehicle (veh)-Nrx mice (C) Representative images showing Fizz1 immunofluorescence (FITC) in macrophages and bronchial epithelial cells corresponding to lung sections of veh and M2_regs_-treated Nrmx or Hpx mice; cell nuclei are stained with DAPI; arrows show Fizz1-stained macrophages in airspaces; scale bar = 50μm. (D) Quantification of *Il6* and *Ccl17* mRNA and Fizz1protein levels determined as mean fluorescence intensity (MFI) normalized for cell number (DAPI stain), in bronchoalveolar (BAL) cell pellet from mice housed in Nrmx or Hpx for 3 days. (E) Representative BAL cytospins showing Fizz1 immunofluorescence staining in veh- and M2_regs_-treated Nrmx or Hpx mice; scale bar = 12.5μm. Mean ± SEM, n=3-5 per group, *p <0.05, **p <0.01, ***p <0.001, ****p <0.0001.

### Attenuation of inflammation occurs through early reduction of monocytic recruitment, accumulation of CD68 macrophages and complement activation

Beyond increased expression of inflammatory cytokines, monocyte/macrophage recruitment and perivascular accumulation are major immunoregulatory components contributing to human PAH and preclinical PH. We next investigated the effect of M2_regs_ administration on pulmonary myeloid cell subsets after 2 days of hypoxia exposure. Flow cytometry analysis from total lung allowed us to identify alveolar and interstitial macrophages as being CD11c^high^, MHCII^+^, CD64^+^, CD11b^-^ and CD11b^high^, MHCII^+^, CD64^+^, CD11c^low^, respectively. Non-classical monocytes are defined as CD11b^high^, MHCII^-^, CD64^+/-^, Ly6C^low^, while classical monocytes as expressing CD11b^high^, MHCII^-^, CD64^+/-^, Ly6C^high^ [27]. Cell suspensions prepared from lung digests were gated according to the strategy depicted in Fig. 5A. Flow cytometry of lungs obtained from Hpx mice showed an increased proportion of alveolar macrophages compared to Nrmx counterparts that was not ameliorated by M2_reg_ treatment. In addition, frequencies of interstitial macrophages were increased by hypoxia, but their numbers remain comparable in both treated and untreated groups. Recruitment of classical monocytes was induced while that of non-classical monocytes was decreased after 2 days of Hpx exposure, but this effect on monocytic frequencies was attenuated after M2_reg_ administration (Fig. 5B).

**Figure 5.**
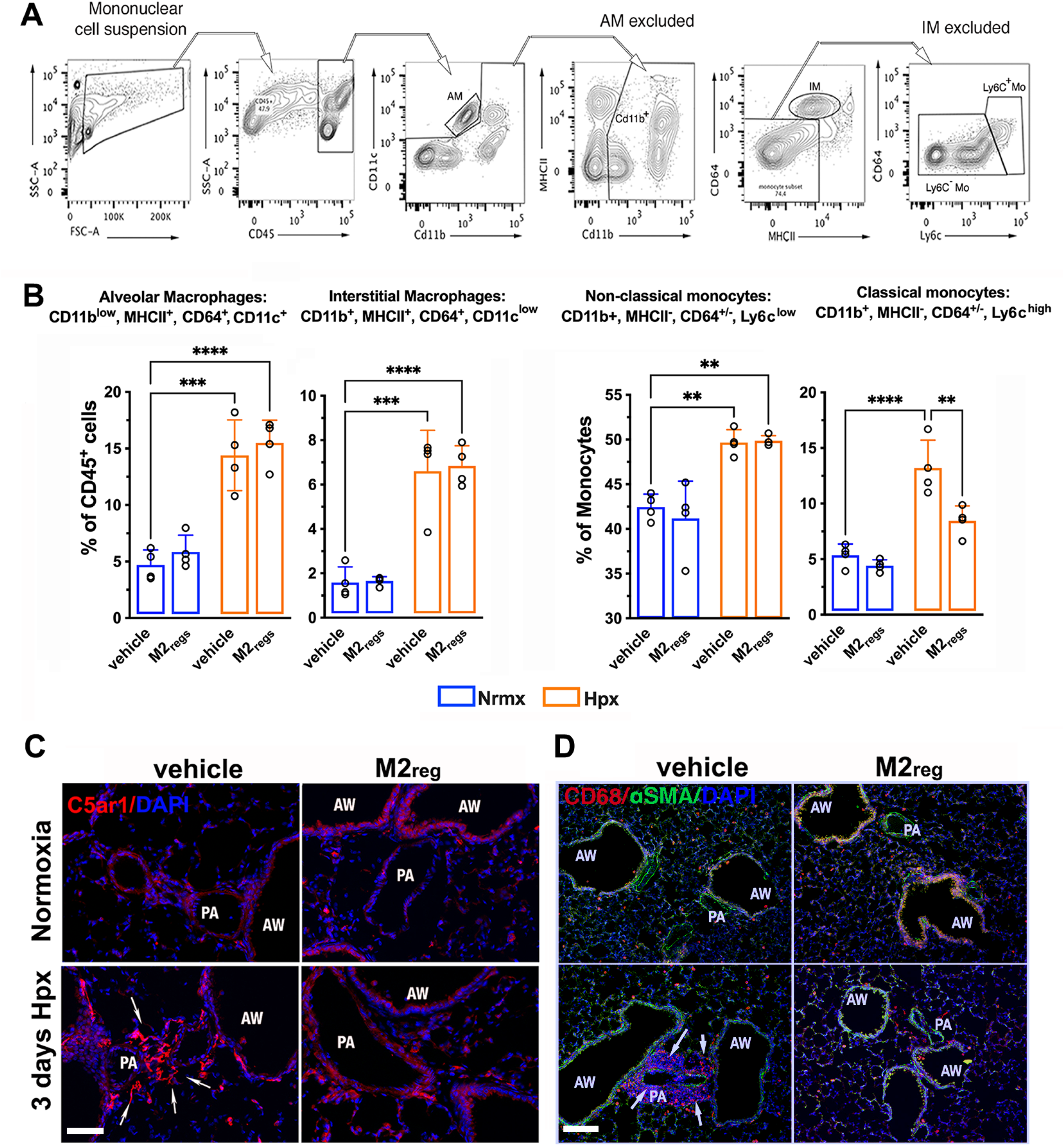
M2_regs_ attenuate monocytic recruitment and perivascular macrophage accumulation. (A) Representative flow cytometry contour plots showing the gating strategy used to identify monocytes and macrophages in lung mononuclear cell suspensions from 2-day housed normoxia (Nrmx) or hypoxia (Hpx) mice having received media (vehicle) or M2_regs_. (B) Histograms showing frequencies (as percentages of the parent cell) of alveolar (CD11b^low^, MHCII^+^, CD64^+^, CD11c^+^) and interstitial (CD11b^+^, MHCII^+^, CD64^+^, CDl1c^low^) macrophages, and non-classical (CD11b^+^, MHCII^-^, CD64+/-, Ly6c^low^) and classical (CD11b^+^, MHCII^-^, CD64+/-, Ly6c^high^) monocytes in each experimental group. (C) Representative images of lung complement C5a receptor 1 (C5ar1) perivascular immunostaining in vehicle and M2_reg_-treated mice after 3 days of Hpx exposure. Arrows show increased C5ar1staining in areas surrounding airway (AW)-associated pulmonary arteries (PA) from vehicle-treated tissue sections, compared to M2_reg_-treated lungs; Scale bar = 50μm. (D) Immunolocalization of CD68^+^ macrophages is shown in PAs (arrows) of vehicle treated murine lungs exposed to 3-day Hpx and absent in lung sections from M2_reg-_treated counterparts. Scale bar = 100μm. Mean ± SEM, n=4 per group, **p <0.01, ***p <0.001, ****p <0.0001.

As complement activation and perivascular macrophage recruitment are hallmarks of the pathobiological mechanisms mediating inflammation and remodeling in experimental hypoxic PH [7], we examined whether perivascular accumulation of CD68 macrophages is attenuated by M2_reg_administration and whether this accumulation is accompanied by increased complement activation in Hpx lungs. Three days after hypoxia exposure, lung sections from vehicle-treated mice showed increased expression of complement C5a Receptor 1(C5ar1) in airway-associated pulmonary arteries while staining of this receptor in M2_reg_-treated mice was comparable to the one observed in Nrmx-exposed mice (Fig 5C). In addition, macrophage accumulation was detected in the lungs of 3 days Hpx-exposed mice, and this increase was associated with perivascular compartments (Fig 5D). Mice that received M2_regs_ had reduced numbers of CD68^+^ macrophages in pulmonary arteries, suggesting that complement activation initiated by hypoxia leads to selective and progressive macrophage accumulation in the lung and treatment with M2_regs_ inhibits this response.

Early macrophage activation and induction of inflammatory markers are required for hypoxic-induced vascular injury and establishment of PH [14], and M2_reg_ protection seems to curtail these early events. To examine M2_reg_ effect at a later time point, after disease has developed, we adoptively transferred M2_regs_ to mice 14 days after the initiation of Hpx and evaluated PH development and lung macrophage accumulation 28 days post-Hpx (Figure 6A). As expected, 4 week-Hpx induced macrophage accumulation and RVH development and early M2_reg_ administration mitigated those effects. However, late M2_reg_ treatment (d14) did not prevent CD68^+^ cell increase (Figure 6B) nor ameliorate Fulton’s Index elevation (Figure 6C) when compared to M2_reg_ administration at the beginning of the exposure (d0), revealing a critical role of M2_regs_ during the initial inflammatory phase leading to persistent alveolar and perivascular macrophage accumulation and PH.

**Figure 6.**
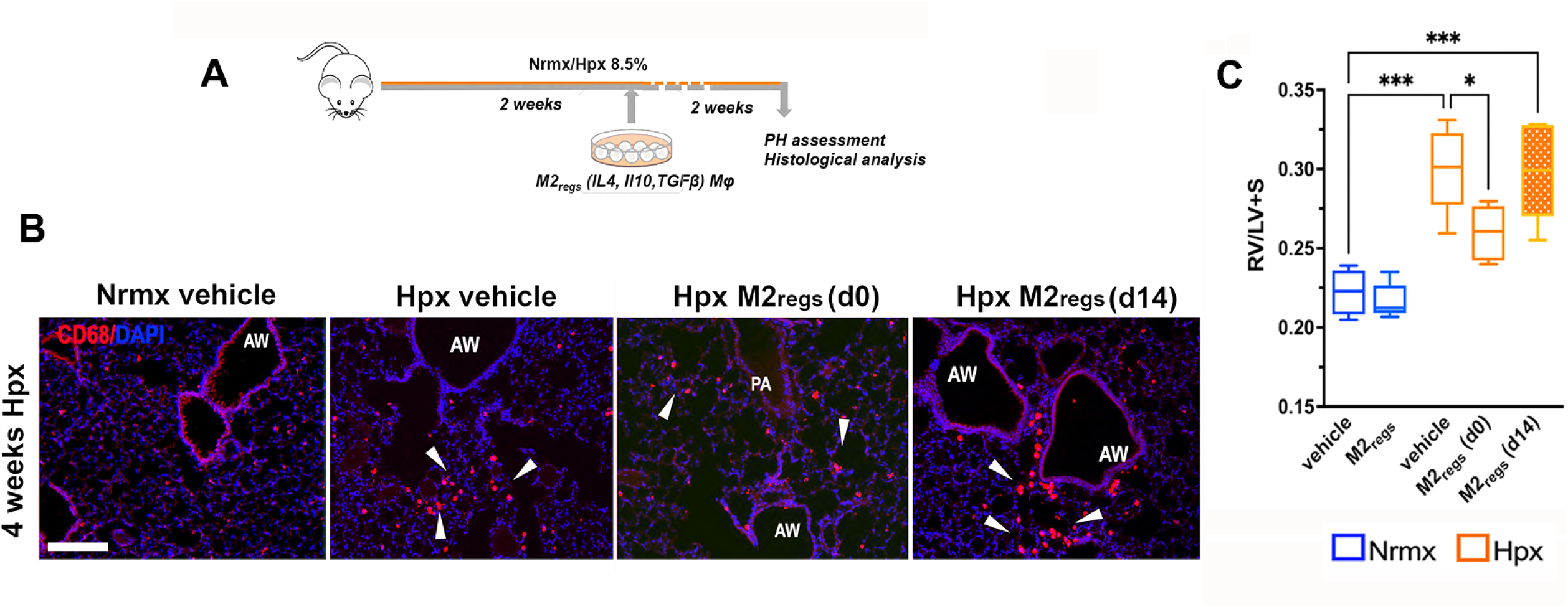
Late M2_reg_ administration fails to reduce perivascular inflammation and PH development. (A) Scheme showing delayed M2_reg_ administration and analysis of PH progression in 4-week housed normoxic (Nrmx) or hypoxic (Hpx) mice. (B) Hpx-induced lung tissue CD68^+^ macrophages accumulation CD86+immunofluorescence was ameliorated in Hpx mice that received M2_regs_ at the beginning (day 0, d0) of the 4-week exposure period but not in mice treated with M2_regs_ at day 14 (d14) after the initiation of Hpx. Arrowheads showing perivascular CD68 immunofluorescence (red); cell nuclei are stained with DAPI. Scale bar = 50μm. (C) Fulton’s Index (RV/LV+S) determined in mice exposed to 4 weeks of Nrmx or Hpx that were left untreated (vehicle) or treated with early (d0) or late (d14) M2_regs_. Mean ± SEM, n=4 per group, *p <0.05, ***p <0.001.

### Administration of M2_regs_ promotes an anti-inflammatory lung milieu

An increase in monocytic infiltration and subsequent elevation of proinflammatory cytokines *Il6*, *Ccl17* and *Fizz1* mRNA expression in the lungs of Hpx-exposed mice suggested that hypoxia induces an inflammatory state that is attenuated by the pulmonary adoptive transfer of regulatory M2_reg_ macrophages. To further investigate the cellular mechanisms responsible for this protection, we used an *ex vivo* assay in which we could assess M2_reg_ potential to modify the local immune microenvironment of the lung. To this end BMDMs, used here as reporter cells, were cultured in the presence of BAL supernatants obtained from 3 days Nrmx or Hpx-exposed mice that were pretreated with vehicle or M2_regs_ (Figure 7A). Proinflammatory cytokine expression, quantified by qPCR showed increased *Il6*, *Ccl17* and *Fizz1* mRNA levels in BMDMs cultured in the presence of hypoxic BAL obtained from vehicle-treated mice. However, when BMDMs were exposed to BAL obtained from Hpx M2_reg_-treated mice, expression of these cytokines was reduced and returned to vehicle-treated Nrmx levels, while *Chi3l3* expression increased compared to those in BMDMs stimulated with BAL obtained from vehicle-treated Hpx mice (Figure 7B).

**Figure 7.**
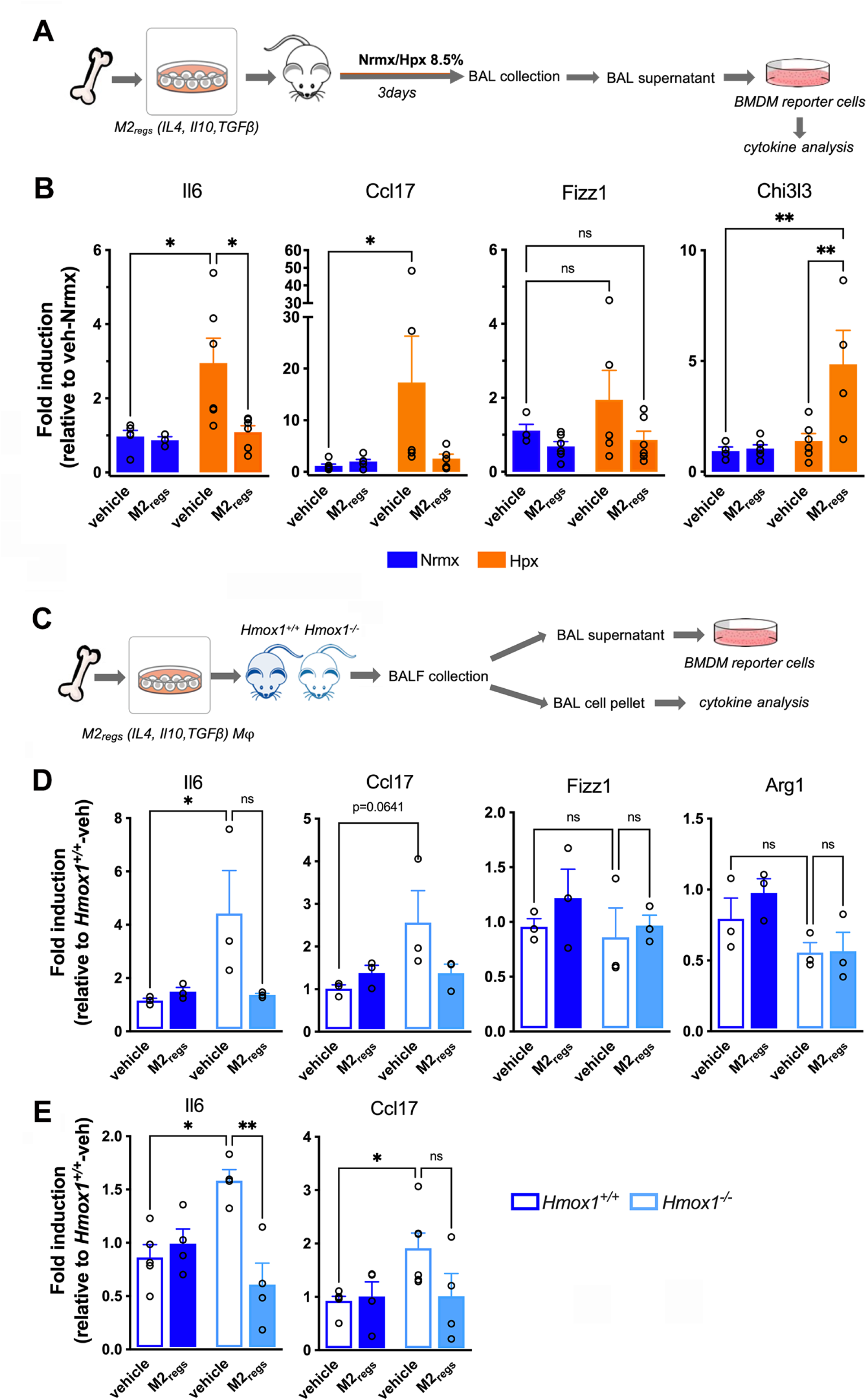
M2_regs_ modulate the lung milieu to suppress pro-inflammatory signals. (A) Scheme showing treatment of bone marrow derived macrophages (BMDMs) with bronchoalveolar (BAL) supernatants obtained from mice receiving media (vehicle) or M2_regs_ and housed in Nrmx or Hpx for three consecutive days. (B) BMDM expression level of cytokines and inflammatory mediators, determined by qPCR, showing normalization (*Il6, Ccl17 and Fizz1*) or augmentation (Chi3l3) by BAL supernatant derived from Hpx M2_regs_-treated mice, compared to supernatants from vehicle-treated counterparts. Expression levels are shown relative to (vehicle) veh-Nrx BAL-treated BMDMs. (C) Scheme showing treatment of reporter BMDMs with BAL supernatants obtained from wild type (*Hmox1^+/+^*) or Hmox1 deficient (*Hmox1^-/-^*) mice treated with vehicle or M2_regs_. (D) *Hmox*1^-/-^ BAL supernatants induced the expression of *Il6* and *Ccl17* mRNA in BMDMs compared to *Hmox1^+/+^* BAL, with little effect on Fizz1 and Arg1 gene expression. (E) *Il6* and *Ccl17* mRNA expression in cell pellets from BAL fluid from the same experimental groups. *p<0.05; **p<0.01.

To further investigate the regulatory effect of M2_regs_ on the lung environment, we took advantage of the inflammatory phenotype induced by heme oxygenase-1 (*Hmox1*) deficiency. In *Hmox1* deficient macrophages, increased NF-kB activation, and elevated expression of cytokines *Il6*, *Ccl17* and *Ccl2* is elevated compared to wild type macrophages [24]. We assessed the potential of wild type (*Hmox1^+/+^*) and deficient (*Hmox1^-/-^*) BAL supernatant to induce cytokine production in cultured BMDMs using our *ex vivo* assay (Figure 7 C). BMDMs exposed to *Hmox1^-/-^*BAL supernatants from vehicle-treated mice showed increased mRNA expression of *Il6* and *Ccl17* (Figure 7D) with little effect on Fizz1 and Arg1 expression. However, the expression of these cytokines was suppressed by BAL supernatants obtained from *Hmox1^-/-^*-M2_reg_ treated-mice suggesting that M2_regs_ dampen the inflammatory lung milieu of Hmox1 deficiency. In addition, induced expression of *Il6* and *Ccl17* mRNA in BAL cell pellet mirrors the one achieved with the corresponding BAL supernatant (Figures 7E) suggesting that these cytokines may originate from pulmonary alveolar macrophages and that adoptively transferred M2_regs_ may participate, at least in part, in the regulation of the local microenvironment.

## DISCUSSION

In this study we show that M2_reg_ administration attenuates monocytic recruitment and perivascular inflammation that occurs with hypoxic exposure, and reduces early lung and alveolar macrophage inflammation, events leading to PH development. Furthermore, our results indicate that M2_regs_ adopt an anti-inflammatory phenotype that can be partially maintained *in vivo* and can modify the inflammatory signature of the immune microenvironment caused by hypoxia upon their recruitment to the airspace.

Therapeutic interventions for clinical PAH are limited to those targeting specific cytokines and inflammatory mediators that limit vasodilatation and diminish RV afterload [28]. Elevated circulating levels of the proinflammatory cytokines TNF⍺ and IL6 predict poor outcomes in idiopathic and familial patients with PAH and pharmacological targeting of their receptors ameliorates experimental PAH [16, 29]. However, these targeting approaches often fail due to the complexity of cell types and signaling pathways involved in the pathogenesis of PH. Reshaping pulmonary monocyte-macrophage recruitment and macrophage activation has been the goal of recently proposed anti-inflammatory treatments in experimental PH [30–32]. Efforts to reduce their mobilization into the lung using genetic or pharmacological approaches have resulted in reduced vascular remodeling and partial RVH normalization [8, 12]. In line with these reports, the beneficial effects of the pulmonary administration of M2_regs_ in our study appear to be mediated, at least in part, by reducing the recruitment of classical Ly6C^+^monocytes in the lung, while maintaining the proportion of non-classical Ly6C^-^ monocytes to values comparable to those of Nrmx mice. In this regard, adoptive macrophage transfer approaches seem advantageous against depleting strategies as the latter may lead to unwanted removal of beneficial macrophages at specific times and compartments. For example, non-classical monocytes are integral cellular components of the lung vasculature and enter the tissue at steady state to become interstitial macrophages [33]. Recruited monocyte-derived macrophages in the alveolar space suppress alveolar macrophage inflammation, neutrophil accumulation, and loss of epithelial barrier function in a model of acute lung injury [34] and are critical for the resolution of bleomycin-induced lung injury during the fibrotic period [35] and the regression of vascular remodeling induced by sub-chronic exposure to hemoglobin and hypoxia [36].

Macrophage activation and repolarization during the initial period of hypoxia are further events contributing to lung injury and PH development[14, 16, 37]. Activation of resident alveolar macrophages seems to initiate local inflammation and monocyte migration into the alveolar and interstitial compartments [38] and contribute to the chronic phase of alveolar hypoxia [39]. In our study, the initial induction of *Il6*, *Ccl17* and *Fizz1* expression in the lung is paralleled by similar cytokine elevations in alveolar macrophages suggesting that these cells are in part responsible for their production. M2_reg_ administration attenuates the increased expression of these cytokines while enhancing the expression of Arg1, indicating that they may act locally to improve alveolar macrophage function and halt monocyte migration. Progression of hypoxia-induced pulmonary inflammation occurs with increased accumulation of interstitial macrophages, in agreement with previous reports [8, 15]. Of note, the number of CD68^+^ perivascular macrophages increase progressively with the hypoxic exposure and adoptive transfer of M2_regs_ reduced their accumulation while delayed administration of M2_regs_ did not, confirming the importance of early alveolar macrophage activation for the continued recruitment and expansion of interstitial monocyte-derived macrophages. As Frid et al., recently demonstrated [7], complement activation greatly contributes to vascular remodeling. Attenuation of medial vessel wall thickening, muscularization of small arterioles and reduction in hemodynamic function in our study may at least be partially due to complement inhibition by M2_regs_ treatment.

Our results show that M2_regs_ protect the lung from hypoxic injury and PH development and that these cells have the potential to modify the immune microenvironment of the alveolar space during early hypoxic inflammation. Classical pro-inflammatory cytokines such as IlL6, Ccl17 and Tnf⍺ are suppressed by M2_regs_. This inhibitory effect by M2_regs_ may prevent the early inflammatory response to hypoxia. On the other hand, M2_regs_ dramatically induce Arg1 which is an important inhibitory enzyme for the nitric oxide pathway, but also critical for resolving inflammation through converting arginine to ornithine and promoting efferocytosis with resolution of injury [40]. The actions of Chi3l3 (Chitinase 3-like-3/Ym1) on immune modulation, also induced by M2_regs_ in our study, are not fully understood in regard to lung injury. A recent report, however, demonstrated a protective role of Chi3l3 produced by brain macrophages and microglia in an experimental model of neuroinflammation [41]. Combined, inhibition of early inflammatory cell recruitment in the hypoxic lung and enhanced resolution of inflammation may be key mechanisms of the protection conferred by M2_reg_ adoptive transfer. An open question is whether M2_regs_ suppress macrophage activation and monocyte recruitment directly through factors and mediators released to the alveolar niche, e.g., TGFβ and IL10, microvesicles, etc., or indirectly through interaction with resident alveolar macrophages similarly to the way recruited monocyte-derived macrophages suppress alveolar macrophage inflammation in acute lung injury[34].

Anti-inflammatory M2_regs_ have been implicated in the suppression of reactive T-cells in a diabetes mouse model [22] and generation of Treg [42] in an induced model of colitis, and both types of T cells have been involved in the initiation and progression of PH[43, 44]. Adoptive transfer experiments have demonstrated that macrophages show a remarkable plasticity when they are transplanted into another tissue environment [45, 46], acquiring phenotypic characteristics of resident cells. In our study, transplanted bone marrow derived M2_regs_ are retained in the lung and are reshaped into an alveolar macrophage phenotype that partially acquires an inflammatory nature over time from the hypoxic alveolar microenvironment, in agreement with previous transcriptional studies performed in the aging lung [47] indicating that changes in their phenotype are not dependent on intrinsic signals but are modified by environmental cues.

This study has several limitations. First, we have not made attempts to deplete the alveolar niche with the use of intratracheal liposomal clodronate and therefore the contribution of tissue resident cells to the anti-inflammatory effects of M2_regs_ has not been completely addressed. In addition, M2_reg_ engraftment is poor and probably a reflection of niche availability and may have limited the beneficial effects of the adoptive transfer. Moreover, one single M2_reg_ administration could have delayed but not prevented the peak of hypoxia as we have previously demonstrated using inducible *Hmox1* overexpression in the hypoxic lung [14] Future experiments will address if PH prevention by M2_regs_ requires suppression of macrophage activation and accumulation of the entire period of hypoxia-induced inflammation.

Considering the dynamic changes associated with hypoxia-induced PH, M2_regs_ adoptive transfer has the potential to advance the resolution of inflammation while blocking its initiation and continuation and should be further explored as a therapeutic approach in treating this condition.

## ACKNOWLEDGMENTS

The authors thank Dr. Eleni Delavogia (Boston Children’s Hospital, Harvard Medical School, Boston, MA) and Mr. John Cortinas (Boston Children’s Hospital) for their scientific discussions. The authors also thank Suzan Lazo, director of Dana Farber Flow Cytometry Cores (Dana Farber Cancer Institute) for providing training and assistance with the acquisition of flow cytometry experiments.

## Author Contributions

A.F.-G., and A.M. participated in study design and execution, data collection, analysis, and manuscript writing. J.N. and K.Z. participated in study execution and analysis. S.V. performed right ventricular systolic pressure measurements and data analysis. G.W. and M.R. contributed with collection, analysis, and interpretation of data. A.G. provided data analysis and interpretation. X.L. contributed with animal technical support. S.A.M. and S.K. contributed to study design, supervision of study execution, analysis, financial support, manuscript writing, and the editing and approval of the final article.

Supported by NIH 5R01HL055454, RO1HL146128 (to SK)

---

**Supplemental Figure 1.** Gating strategy used to characterize the phenotype of untreated (M0), LPSγ/IFNγ (M1) and IL4/IL10/TGFβ (M2_reg_) bone marrow derived macrophages (BMDMs) prior to their *in vivo* adoptive transfer.

**Supplemental Figure 2.** Gating strategy used to characterize *ex vivo* DiD-labeled M0 and M2_regs_ obtained from adoptively transferred mice exposed to 4 weeks of normoxia (Nrmx) or hypoxia (Hpx).

**Supplemental Figure 3.** Adoptively transferred M2_regs_ remain in the alveolar space long after *in vivo* delivery. Detection of DiI-labeled M2r_egs_ in lung sections after 4 weeks of oropharyngeal (ET) or intranasal (IN) administration and loss of DiI-labeling in lungs undergoing bronchoalveolar (BAL) lavage. (B) Cytospins prepared from media (vehicle) and M2_reg_-treated mice exposed to normoxia (Nrmx) or hypoxia (Hpx) for 3 days. Cell nuclei are stained with DAPI.

